# Survivin prevents the Polycomb Repressor Complex 2 from methylating Histone 3 lysine 27

**DOI:** 10.1101/2022.04.27.489567

**Authors:** Maja Jensen, María-José García-Bonete, Atsarina Larasati Anindya, Karin Andersson, Malin C. Erlandsson, Nina Oparina, Ulrika Brath, Venkataragavan Chandrasekaran, Bokarewa I. Maria, Gergely Katona

**Affiliations:** Department of Chemistry and Molecular Biology, Faculty of Science, University of Gothenburg, Sweden; Department of Medical Biochemistry and Cell Biology, Institute of Biomedicine, University of Gothenburg, Gothenburg SE-405 30, Sweden; Department of Rheumatology and Inflammation Research, Institute of Medicine, University of Gothenburg, Box 480, 40530 Gothenburg, Sweden; Department of Chemistry and Molecular Biology and the Swedish NMR Centre, University of Gothenburg, SE-412 96 Gothenburg, Sweden; Rheumatology Clinic, Sahlgrenska University Hospital, Gröna stråket 16, 41346 Gothenburg, Sweden

## Abstract

Survivin is a small protein that belongs to the inhibitor of apoptosis protein family and participates in cell division and apoptosis. It was actively studied in human cancers, inflammatory diseases and in autoimmune diseases. Here, we reveal that survivin takes part in epigenetic gene silencing by interaction with the polycomb repressive complex 2 (PRC2). PRC2 silences gene expression through tri-methylation of lysine 27 on histone 3 (H3K27). We detected differential expression of PRC2 core subunits in CD4+T cells with different survivin expression. ChIP-seq experiments indicated that survivin binds chromatin that overlap with the regions occupied by PRC2. ChIP-seq of H3K27 in CD4+T cells indicate that inhibition of survivin leads to a substantial increase in H3K27 tri-methylation by PRC2 in contrast to other histone modifications, which lends support to that survivin prevents PRC2 from functioning. Survivin binds peptides derived from PRC2 subunits EZH2, EED, SUZ12 and JARID2 in a peptide microarray that cover intersubunit interfaces, catalytic residues, and present binding sites for substrates, DNA, and regulatory proteins. Amino acid composition of the peptides has substantial predictive power for survivin interaction in the peptide microarray as determined by multilayer perceptron classification analysis. NMR experiments with ^15^N labelled survivin indicate that peptide colocalization does not entirely depend on binding mediated by short range interactions. These results indicate that survivin interacts with PRC2, preventing the methylation of H3K27 and specific gene silencing. This has transcriptional consequences and specific gene silencing.

## Introduction

For the human body to develop and maintain cell homeostasis a tight regulation of cellular processes, such as apoptosis and cell division is necessary. Survivin is one of the proteins that play an important role in these processes. With a length of 142 amino acids and a molecular weight of 16.5 kDa, survivin is the smallest member of the IAP family. In contrast to the other members of the IAP family, survivin has a single baculovirus-inhibitor of apoptosis repeat (BIR) which is connected to the α-helical C-terminus. ^1,2^ The expression level and location of survivin in the cell is highly dependent on the phase of the cell cycle. During cell division, survivin takes part in the chromosomal passenger complex (CPC) together with Aurora B kinase, INCENP and borealin.^3,4^ When not part of the CPC, during interphase, survivin is in the cytoplasm and prevents apoptosis. During apoptotic signaling Smac/DIABLO is released from the mitochondria and prevents survivin from inhibiting apoptosis. A relatively unknown function of survivin is its role as a transcription factor or cofactor in gene regulation. A more systematic search of survivin binding sites in the chromatin is necessary to strengthen this emerging view.^5^ In order to map potential association with chromatin we performed deep sequencing of DNA regions bound by survivin in CD4+ T-cells.

We found that the precipitated regions were strongly associated with chromatin locations of subunits associated with PRC2. PRC2 is a histone methyltransferase whose main function is to methylate lysine 27 on histone 3 (H3K27), thus regulating gene expression by epigenetic modification. ^6^ When H3K27 is trimethylated the genes are maintained in a repressed state, preserving the identity of the cell. ^7^ PRC2 is intensively studied because of its role in various types of cancer and its ability to restore pluripotency. ^8–10^ The structure of its core subunits is well characterized. The core consists of four subunits; enhancer of zeste homologue 2 (EZH2), embryonic ectoderm development (EED), suppressor of zeste 12 (SUZ12) and retinoblastoma binding protein 46/48 (RBAP46/RBAP48). EZH2 contains the enzymatic component of the PRC2, this is where the trimethylation of H3K27 is taking place. The seven WD40-repeat containing protein EED is the smallest component of the complex. EED has an aromatic cage, which consists of four aromatic side chains, Phe-97, Tyr-148, Trp-364 and Tyr-365. This motif binds the already triple methylated H3K27, stimulating activity of the complex and autocatalysing the methylation of other histone subunits by EZH2.^11^ SUZ12 encompasses the other subunits contributing to the stability of the complex and keeping them together.^12^ Jumonji and AT-rich interaction containing domain 2 (JARID2) and adipocyte enhancer-binding protein 2 (AEBP2) have been reported as important cofactors of PRC2. JARID2 is mimicking a methylated H3 tail, which triggers PRC2 activity. AEBP2 on the other hand is mimicking an unmodified H3 tail. ^12^

EZH2 is the major mediator of PRC2 function in gene regulation in cells. Proper function of EZH2 is critical for normal cell function, while aberrant expression has been connected to proliferation, cell cycle progression and oncogenesis. ^8-10^ Recent studies revealed the importance of EZH2 for homeostasis of hematopoetic stem cells, thymopoesis and lymphopoesis.^13,14^ EZH2 deficiency in T cells seriously affects differentiation ability and function of T cells. ^15-17^ This makes it an attractive intervention target in various forms of cancer and autoimmune conditions. ^18-20^

Here, we studied the role of survivin as a transcription factor in association with chromatin and identified its interaction with PRC2. Specifically, we studied the direct interaction of PRC2 with survivin in peptide microarrays, biolayer interferometry (BLI), and NMR spectroscopy. Machine learning analysis indicated that the interaction between survivin and PRC2 derived peptides is not primarily governed by the sequence of the peptides, but their amino acid/atomic composition. NMR titration experiments revealed that this predictable tethering association does not likely involve binding changes on the surface of survivin. We argue that to explain the location and activity of the PRC2 complex together with survivin, tethering type of interactions are more important than binding type of interactions.

## Results and Discussion

It is known that survivin is binding nucleosomes through phosphorylated Thr-3 of histone H3^21^. Although PRC2 also targets the N-terminal tail of H3, the two actors may not be present at the same time. We explored if H3K27 histone modifications involved in epigenetic regulation colocalize with survivin using ChiP-seq analysis of the DNA material precipitated with antibodies against survivin, H3K27me3 and H3K27ac.

Figure 1A shows a heatmap of survivin peaks H3K27me3 peaks of human primary CD4+ T cells obtained in ChIP-seq experiments. Annotation of the survivin and H3K27me3 peaks to the genome revealed that a significant proportion of these peaks binds to the same DNA regions (Figure1B). In addition to a substantial overlap, the colocalized survivin peaks and histone H3K27 modifications were associated with high peak score (Figure1C). The combination of histone H3 modifications is associated with the highest score of survivin peaks, which argues for the affinity of survivin and histone H3K27 marks to those regions in material of the primary CD4+T cells (Figure 1E).

**Figure 1.**
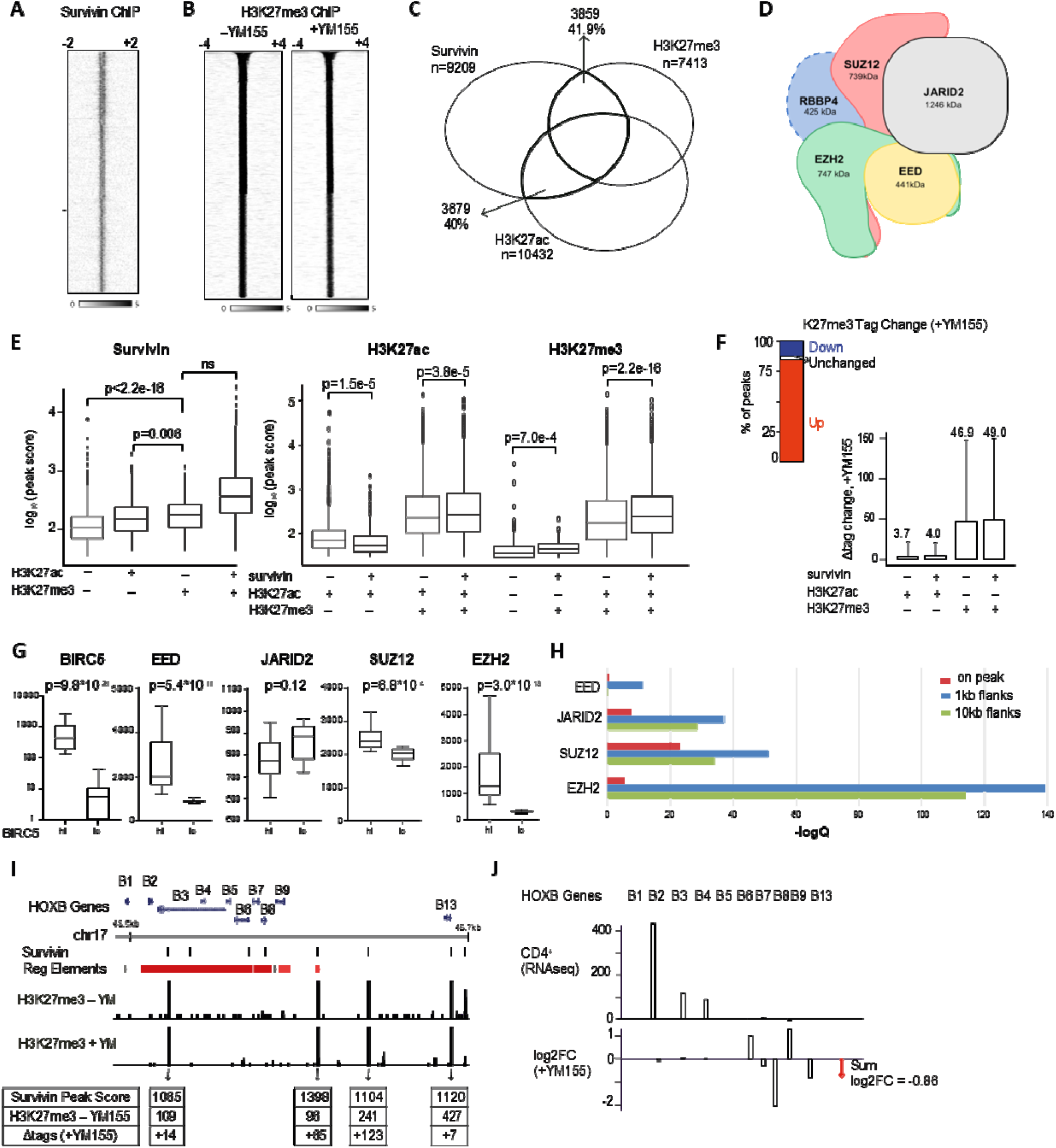
Survivin colocalizes with histone H3K27 marks and controls PRC2 function in human CD4 cells. Heatmap depicts survivin (A) and histone H3K27me3 (B) ChIPseq peaks in human primary CD4 cells, before and after YM155 treatment. (C) Venn diagram of the genomic co-localization between survivin with histone H3K27 ChIPseq peaks in CD4 cells. The peaks overlapping by at least 1bp were considered co-localized. (D) A cartoon of the PRC2 complex. (E) Peak scores of survivin and histone H3K27 epigenetic marks at co-localization sites. Median, IQR and range indicated. Kolmogorov-Smirnov (KS) test p-values are indicated. (F) Proportion change in H3K27me3 peaks after YM155 treatment. The column plot presents the tag count change in H3K27me3 and H3K27ac peaks. Mean and SD indicated. (G) Expression of the PRC2 genes in CD4+ cells (n=24) with high and low survivin/BIRC5, by RNA-seq. Samples were split on the median level of BIRC5 and analyzed using DESeq2. Median with IQR and range, normalized p-values indicated. (H) Genomic colocalization of survivin ChIP peaks with the proteins of PRC2 complex. Regions around Survivin peaks with 0, 1 and 10 kb flanks were investigated. ChIP-seq data of EZH2, SUZ12, EED and JARID2 were obtained through the ReMap database. (I) The cluster of HOX-B genes in the chromosome 17 is transcriptionally controlled by PRC2. Location of Survivin and H3K27me3 peaks in the genome bowser is indicated. *Cis*-Regulatory elements were retrieved from the GeneHancer database. Increase in H3K27me3 peaks co-localized with survivin after YM155 treatment. (J) Expression of the HOX-B genes in human CD4^+^ cells and the expression change after YM155 treatment, by RNAseq. Red arrow on the right indicates the cumulative reduction in the expression of HOX-B genes after Survivin inhibition.

In the next step, we investigated if survivin ChIP-seq peak locations overlap with deposition of the core PRC2 subunits EZH2, EED, SUZ12 and JARID2 (Figure D) supporting a potential concerted action towards H3K27 residue. The integration of survivin peaks with human ChIP-seq mega data for individual PRC2 subunits available through ReMap database (REF) demonstrated significant degree of co-localization with all PRC2 subunits (Figure 1H). Specifically, the proteins of the PRC2 complex were identified within top 10% of the probable survivin partners on the chromatin and dominated by association of survivin peaks with the catalytic EZH2 subunit. The strongest significance was obtained when colocalization was analyzed in the 1k base interval in contrast to direct overlap or wider 10k base window. This indicates that survivin may not directly compete for the DNA region corresponding to the modified nucleosomes, but it acts within a larger protein complex. To pave a functional link the two systems, we utilized 28 RNA-seq datasets of primary CD4+ T cells to study transcription of PRC2 complex. Figure 1G shows a strong association of the core PRC2 subunit expression with the expression level of survivin and presents an additional piece of evidence for concomitant action of survivin and PRC2.

To further investigate the functional role of survivin in modification of histone H3K27 residue, we performed ChIP-seq studies with and without sepantronium bromide (YM155) present, which inhibits survivin transcription^22^, but not expression of cIAP2, XIAP, Bcl-2, Bcl-XL, Bad^22^, cIAP1, p53 or STAT3^23^. The YM155 treatment appear to prevent transcription factor Sp1 to access survivin promoter thus reducing survivin transcription.^24^ YM155 also targets the transcription factor ILF/NF110^25^ it downregulates ID1 expression but stimulates CYLD and FOXO1.^26^ Nuclear translocation of NF-kB heterodimeres (p65/p50) was also disturbed after YM155 treatment.^27^ Despite of the above mentioned side effects of YM155, survivin is suppressed specifically, independently from closely linked proteins.

Comparison of H3K27 peaks revealed that it occurred an increase of H3K27me3 deposition in 85% of the H3K27me3 peaks (Figure 1B, F). A significant increase in tag change in relation to H3K27me3 modification upon survivin inhibition, whereas PRC2 independent H3K27 acetylation remains virtually unaffected (Figure 1F). These results imply that binding of high amounts of survivin in proximity of H3K27 residue may prevent PRC2 from methylating H3K27, which is important for epigenetic silencing of sensitive genes.

To exemplify this function of survivin, we investigated survivin and H3K27me3 peaks in the cluster of Hox-B genes in the chromosome 17. Its transcription is critically dependent on modification of H3K27 residue and PRC2 function^28^. We observed a co-localization of four independent survivin and H3K27me3 peaks, three of which were annotated to the caudal genes between HOX-B8 and HOX-B13 (Figure 1I). Inhibition of survivin with YM155 resulted in an increase in H3K27me3 peaks, by tags (Figure 1I). Transcriptionally, the increase in deposition of inactive H3K27me3 mark led to a change of the HOX-B genes, which was clearly seen in the caudal HOX-B8, -B9 and -B13 genes controlled by those peaks (Figure 1J).

While we cannot exclude that YM155 (or any other survivin suppression techniques for that matter) affect PRC2 activity indirectly, this inhibition evidence should not be viewed independently from the specific colocalization of survivin and PRC2 subunits along the DNA and their affinity to H3 tails.

Based on the *in vivo* results, we were searching for potential direct interactions between survivin and short peptide segments derived from the subunits of PRC2 and other proteins involved histone modifications. We expected that these may be mediated by relatively short-range bonds (hydrogen bonds, ion bridges, van der Waals forces, hydrophobic shape complementarity). The following evidence led to a contradiction with this view and suggested a more intriguing model. Thus, we rationalized the terminology to make the discussion efficient. We will use the expressions “direct interaction” or just “interaction” when we assume that two partners are sufficient for an association, but we do not want to specify the physical mechanism. We will use the word “binding” for describing an association resulting from an initial, random collision between partners and establishing a pattern of short-range physical interactions or “bonds”. We reserve the word “tethering” for specific and predictable interactions resulting in colocalization of partners for which there is no evidence of “binding”, or the interaction “binding” is not plausible due to the great variety of potentially incompatible short range physical interactions. We assume that at least one of the partners must be localized and the partners do not necessarily obey the law of mass action. We will use the word “colocalization” when we simply mean that two or more components occupy a limited shared volume with or without direct interaction.

To shed light on how survivin interferes with the function of PRC2 complex, we designed a peptide microarray of overlapping peptides derived from EZH2, EED, SUZ12 and JARID2. These subunits represent the core of PRC2 and the interaction of survivin with the PRC2 molecular scaffold provides indication about the potentially disruptive role of survivin on the function of PRC2. In Figure 2, we compare the fluorescence signal arising from labeled survivin as a function of position along the polypeptide chain of PRC2 subunits. The raw fluorescence images are displayed in Figure S1. Figure 3 illustrates the microarray fluorescence intensities mapped on the three-dimensional structure of the PRC2 core complex. The entire data set is provided as Supplementary Data 1.

**Figure 2.**
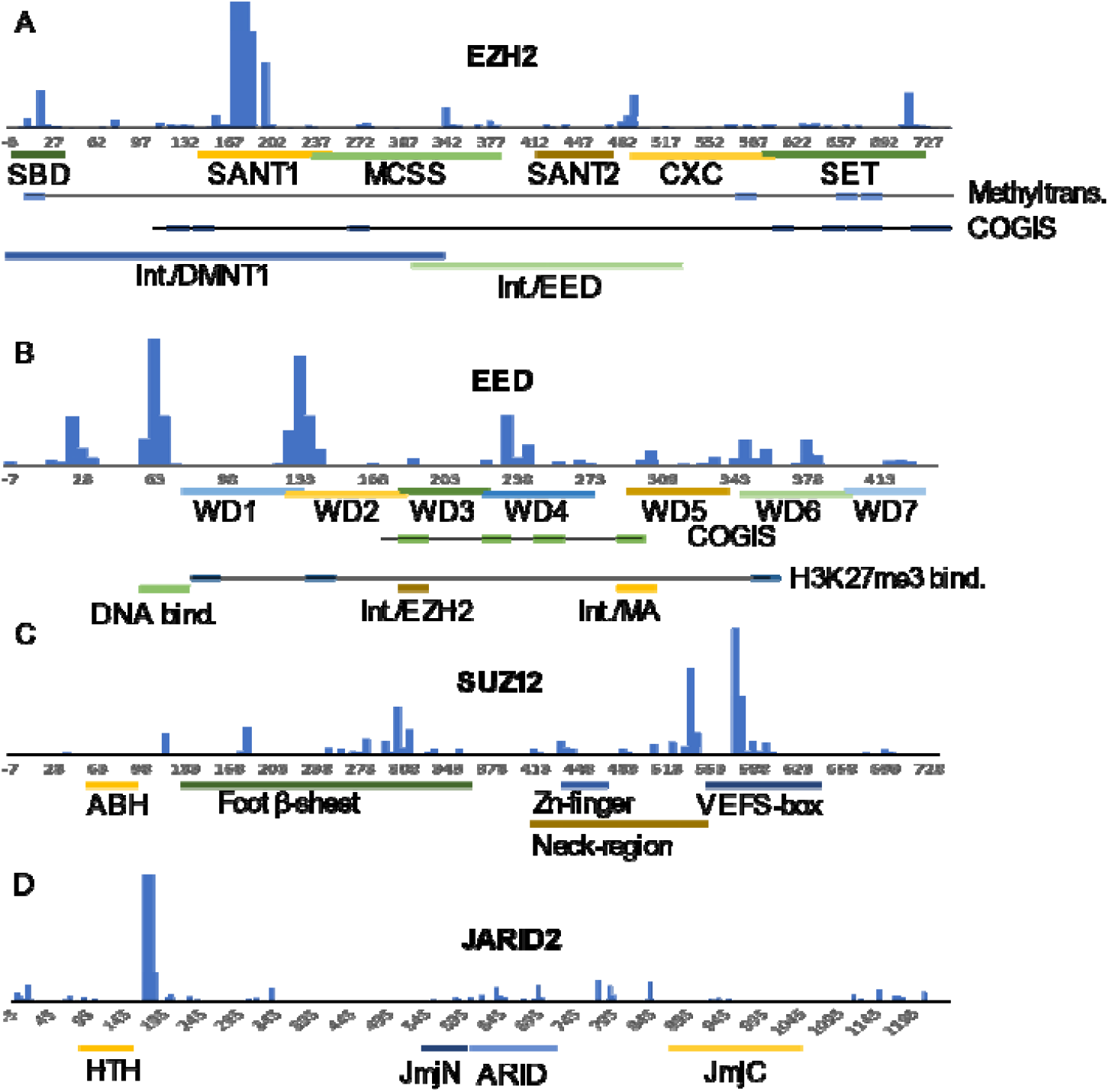
Fluorescence intensity in survivin-peptide microarray of peptides derived from (**A**) EZH2, (**B**) EED, (**C**) SUZ12 and (**D**) JARID2. The residue numbers refer to the first amino acid of the peptides. Negative numbers mean filling amino acid residues that are not part of the natural sequence of the subunit. The annotation bars are inclusive i.e. it is enough to have one amino acid residue from the region on a peptide to be marked as part of the region. Since typically three peptides contain the same amino acid position, the annotation bars overlap even if the described regions do not.

**Figure 3.**
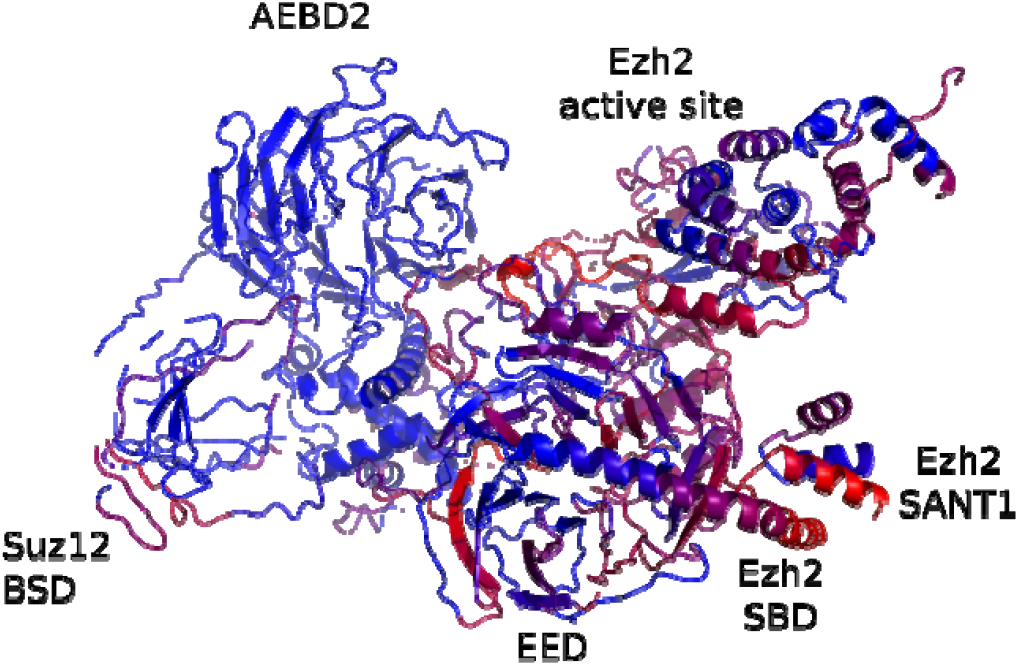
Fluorescence intensity mapped on the 3D structure of the PRC2 core complex (PDB id 6c23)^12^. Blue indicates no interaction or no information about interaction (Aebd2). Progressively brighter red indicates logarithmic magnitude of fluorescence intensity in the peptide microarray experiment. The coloring is inclusive i.e. the residue is associated with the average fluorescence intensity of all the peptides the amino acid residue is present on.

Strongest interaction with EZH2 derived peptides was observed in one continuous region (172-191). The region is located in the SANT1 domain of the enzyme, which is a general binding motif for histone tails and promotes chromatin association.^29^ A strong continuous signal was also observed at peptides in the region 707-736, which included the catalytic residue Tyr-726. Tyr-726 was initially proposed to be the general base for deprotonation of the substrate lysine, but it is likely that the solvent performs this catalytic role.^30,31^ Nevertheless, this is an important part of the active site and when the same region is missing or mutated (Tyr-736, Glu-740) in humans, it results in a specific phenotype described as Weaver’s syndrome, a type of overgrowth disease in the affected patients (Tyr-736, Glu-740).^32^ Lower level of fluorescence was also observed in peptide 627-656 which contains Tyr-641, which is also part of the active site region that brings together the SAM cofactor methyl donor and the H3K27 substrate. ^33^ Y641F mutation in EZH2 results in higher H3K27 tri-methylation activity.^31^ The active site residue Cys-588 is also important for the methyltransferase activity of EZH2 enzyme and peptides containing this residue display moderately high fluorescence. (Figure 2A and supporting online material) Several mutations are associated with overgrowth diseases in EZH2. ^34^ These include the active site residues Y726, Y736 and E740. In addition, survivin binding was observed in a continuous region in the N-terminal direction from the SANT1 region where mutations were associated with overgrowth syndrome (P132 & Y133). ^35^

The SBD domain is located on the N-terminal region of the enzyme, and it is linked to DNA binding. This domain contains residue Ser-21 which is targeted by phosphorylation. Intriguingly, mutation in this position affects the methyltransferase activity of EZH2. ^36^ This region is surprisingly far from the active site in the PRC2 core structure. ^12^ On the other hand, both the entire N-terminal region (residues 1-36) and peptides containing the residues of the active site display high, continuous fluorescence in the survivin interaction assay. The involvement of survivin may interfere or facilitate the connection between these already distant sites and at the same time it may influence the DNA binding properties of EZH2.

In subunit EED, we see repeated (Figure 2B and supporting online material), more evenly distributed regions of high colocalization with survivin including regions 13-37, 58-77, 123-147 and 223-247. The separation of these regions loosely follows the periodicity of the WD repeats, and the regions are concentrated on one side of the doughnut shaped protein. There is a substantial overlap between survivin interacting peptides and the 70-79 region of EED. This region is highly basic and contributes to DNA binding and chromatin positioning. ^37^ Similarly, a mutation at the conserved site R236 is also known as EED-linked Weaver syndrome. ^38,39^ Almost all residues that participate in the binding of the trimethylated lysine are present in the peptides that have high fluorescence signal in the survivin microarray. In addition, H1K26, H3K9 and H3K27 are part of an AXK motif, and the alanine residue of this motif fits into a small pocket on the surface of EED formed by the hydrophobic moieties of Trp-364, Tyr-308 and Cys-324.

In SUZ12 (Figure 2C and supporting online material), the highest affinity peptides (533-607) are concentrated in the C-terminal VEFS-box domain,^40^ but a continuous region with low binding affinity is present between residue range 248-367. Peptides from the zinc finger region (448-471), have relatively small, but noticeable fluorescence signal. Mutation E610V in SUZ12 is known for association with Weaver’s syndrome.^38^ This amino acid residue belongs to the VEFS-box region of the protein.

JARID2 contains a sharply defined interacting region between residues 165-199 and several moderately fluorescent regions. (Figure 2D and supporting online material) The ARID domain is required to direct the PRC2 complex to its target location on chromatin and this domain is abundant with overlapping peptides interacting with survivin. ^41^

The large number of peptides (5395) on the microarray allowed machine learning the characteristics that prompt survivin with interaction. On the microarray approximately 40% of the peptides were associated with larger than zero fluorescence intensity and approximately 20% of the peptides had 1000 or higher fluorescence intensity (Figure S2). Seven peptides with high histidine content were excluded since these reacted strongly with the anti-His-tag antibody directly. Thus, the proportions of interacting and non-interacting peptides are reasonably balanced in this microarray experiment.

Rather than focusing on the sequence of the peptides, we associated the peptides with the abundance of certain atom types in their amino acids (Table S1). Among the many ways to categorize the chemical/positional nature of atoms in the naturally occurring amino acids, the exact choice had limited practical influence on the outcome. Firstly, the peptides were clustered using Ward’s method based on their atom abundance. (Figure 4, Figure S3). The clustering revealed that the similarity in atom type abundance of the peptides is correlated with the fluorescence intensity originating from survivin interaction. On the larger scale, the accumulation of interacting peptides can be observed, which is associated with certain type of atom compositions. The dendogram contains multiple patches of large density of interacting peptides separated by large branches that are not or are very sparsely populated with interacting peptides. This does not appear to be a simple function of the presence or absence of chemical group for example only the presence of large number of carboxyl groups. Rather a specific combination of atom types is required to be present, and the right combination of atoms need to be absent for a peptide to interact with survivin. A zoomed in part of the cluster (Figure S3) illustrates that despite of the different density of interacting peptides in the global dendogram, the suitability of a peptide for interaction depends on the composition in an extremely fine-grained manner. Even peptides with very similar composition can act differently when interacting with survivin. Thus, clustering based on similarity metrics alone may not completely able to group interacting and non-interacting peptides together on the more fine-grained scale. It also means that simple decision trees and trivial arguments are unsuitable for accurately and sensitively predicting whether a peptide is interacting. A trivial argument may simply focus on extremely large net charges or similar extremes of hydrophobicity.

**Figure 4.**
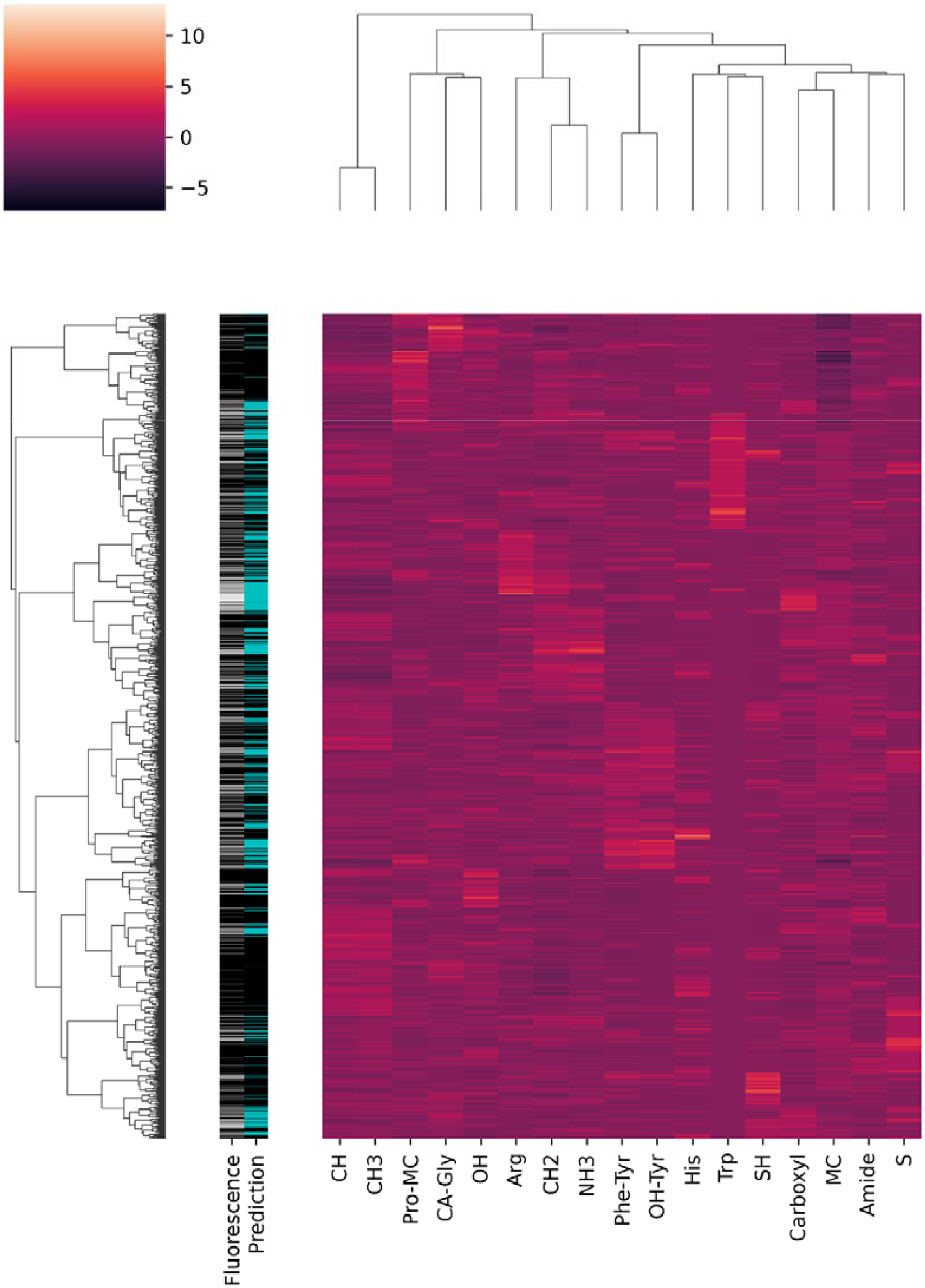
Clustering of peptides based on their atom type abundance (light/dark high/low abundance of atom types, respectively, z-score). The grayscale bar indicates the logarithm of fluorescence intensity of the peptide in the survivin peptide microarray experiment (*black* zero intensity, *white* highest level of intensity). The prediction bar shows the success of the machine learning prediction using the atom type abundance as features (*black* and *cyan* colors mark predicted non-interacting and interacting peptides, respectively).

To go further with the analysis, the peptides were classified into interacting and non-interacting categories, and a neural network was trained on half of the data set to recognize these two classes based on the features of atom composition. The strength of prediction was tested on the remaining half of the data set. The confusion matrix is shown in Table 1. In addition, the predicted interacting peptides are marked with cyan color in Figure 4 and Figure S3. The prediction specificity and sensitivity appeared to be the most successful for the non-interacting peptides providing a solid ground for the negative selection of the peptides, although approximately 2/3 of the interacting peptides were correctly recognized. The density of predictions follows remarkably well the fluorescence signal in Figure 4 and the network often correctly predicts the interacting peptides even among very similar peptides on a more local scale (Figure S3).

**Table 1.**
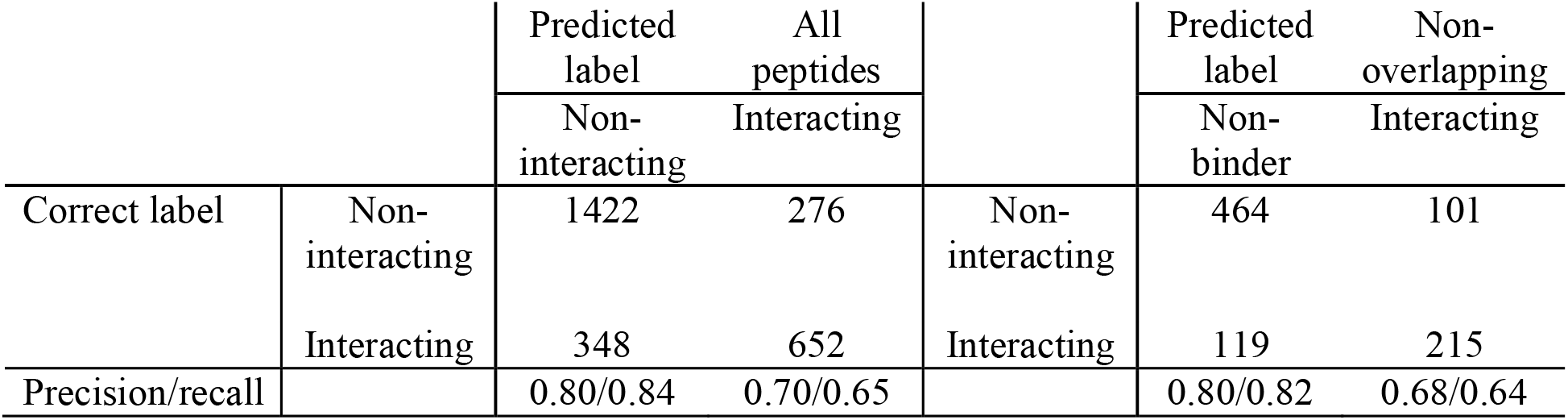
Confusion matrices of predictions

Since the peptides overlap in sequence, the sequence similarity between the test and training set could potentially contaminate the test set. By considering every third peptide only for the test and training set, we eliminated the effect of the overlaps. The relative proportions of elements in the confusion matrix are very similar to relative proportions of elements in the confusion matrix when the classification included every peptide. Therefore, we concluded that the strength of predictions did not originate from training set contaminating the test set. The prediction of interacting peptides did not improve when the interacting peptides with fluorescence intensity below 100 were excluded from the training and testing sets. (Table 2)

**Table 2.** Confusion matrices of predictions with weak interaction partners (0<Fluorescence intensity (FI)<100) excluded from the analysis.

|  |  | Predicted label | All peptides<br>FI >100 or<br>= 0 |  | Predicted label | Non-overlapping<br>FI >100 or<br>= 0 |
| --- | --- | --- | --- | --- | --- | --- |
|  |  | Non-interacting | Interacting |  | Non-interacting | Interacting |
| Correct label | Non-interacting | 1450 | 282 | Non-interacting | 477 | 93 |
|  | Interacting | 297 | 610 | Interacting | 105 | 205 |
| Precision/recall |  | 0.83/0.84 | 0.68/0.67 |  | 0.82/0.84 | 0.69/0.66 |

We also examined the uniqueness of peptide features as described with amino acid composition. Only eight peptides were significantly associated with at least one pair that had identical amino acid composition, again this cannot explain the strength of predictions. The near exclusive uniqueness of peptides is expected from a random model which assumes uniform probability of the 20 amino acids to occur at any position in the 15 amino acid long peptide. The probability of obtaining a specific peptide sequence is 1/20^15^ = 3.1 × 10^−20^, but the abundance of amino acids is not specific to a particular sequence. Since all permutations of the sequence results in identical atom composition, the probability of obtaining a signature composition in a 15 amino acid peptide is 15!/20^15^ = 4 × 10^−8^. In this microarray design, 10 amino acids are constrained between adjacent peptides. Adjacent peptides with identical amino acid composition is still not expected frequently by chance (5!/20^5^ = 4 × 10^−5^) in an array of 5395 peptides. Nevertheless, functional protein sequences are not random. This explains the presence of a minor fraction of peptides that share identical amino acid composition, and such repeated occurrences were exclusively associated with polypeptide chains belonging to the same protein.

Survivin concentration was 1 µg/ml in in the microarray experiment, which suggests that survivin was present in its monomeric form. At this low concentration, there was a clear link between the amino acid composition of the peptide and survivin location. For evaluating the strength of predictions, one should consider that the local torsions and global structure of the peptide was not even considered in this analysis. Moreover, the limit of detection was set to the smallest possible fluorescence signal. At low fluorescence intensity, counting errors could easily result in missing out weak associations. Nevertheless, the rules governing peptide tethering do not necessarily involve specificities of the peptide sequence. They are probably relatively simple since they are inferred from a relatively small training sample set which can work with similar prediction strength as larger data sets. Even the composition of 15 amino acid peptide segments provides a high degree of uniqueness and highly scalable intensity for survivin tethering. Survivin has a remarkable three-dimensional structure that lacks deep binding pockets. Still, survivin can selectively associate with and avoid multiple peptides with signature amino acid compositions that do not require a specific level of net charge or hydrophobicity.

To describe how the interaction affects survivin, ^15^N isotope labeled survivin was titrated while measuring 2D nuclear magnetic resonance spectra (NMR) with PRC2 derived peptides. The peaks associated with assigned protons do not shift substantially during the titration. Figure 5 shows the unclear contrast in NMR chemical shifts in survivin between no EZH2 peptide present and at the maximum concentration of 5 mM. Although not all positions of the peaks are identical, the central part of survivin, where most of the assigned peaks are located, does not substantially change at the highest peptide concentration. The total signal intensity decreases due to the interaction with the peptide. This NMR observation argues against direct binding mediated by short range interactions and shape complementarity as this could introduce chemical shift alterations of surface residues. We cannot completely discount the possibility that the monomeric and dimeric forms of survivin have different affinity to the studied peptides. Since a high survivin concentration is required for NMR spectroscopy, we could not unambiguously connect these results to the observed tethering in the microarray experiment and the observed colocalization and functional links on the cellular level.

**Figure 5.**
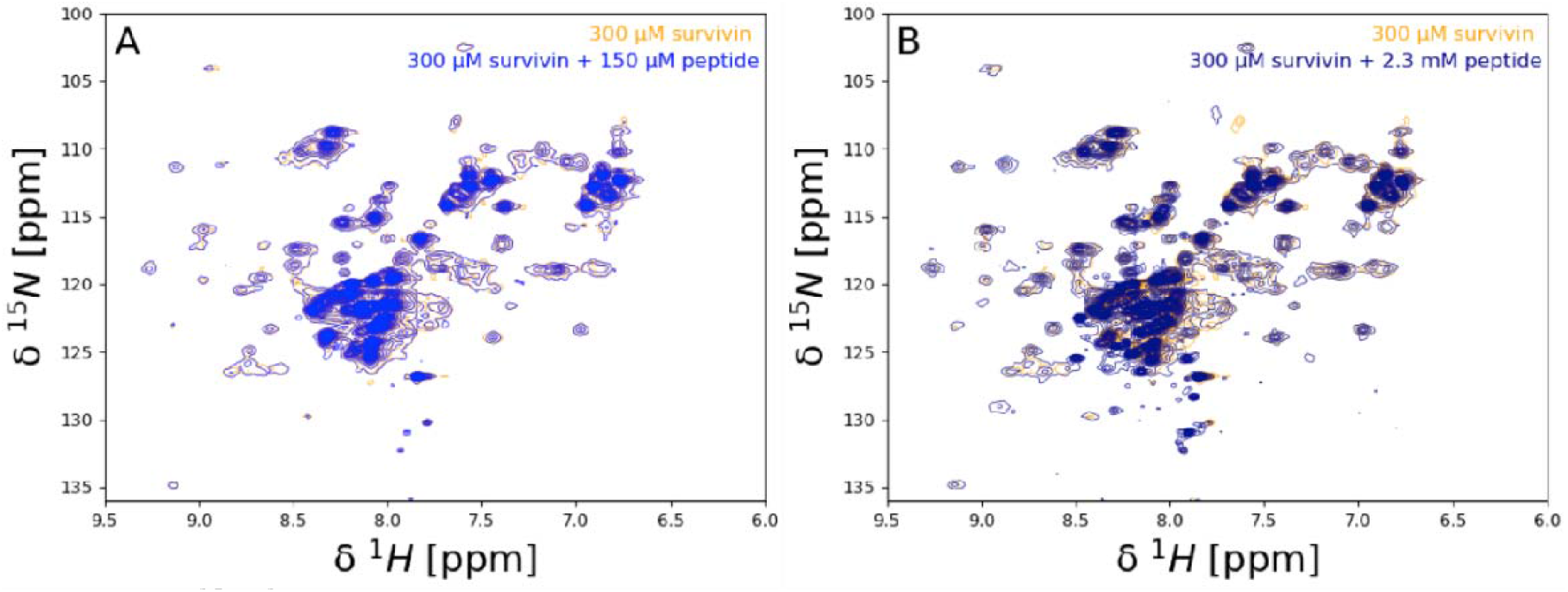
(**A**) Overlay of 2D [^15^N,^1^H] NMR spectra of 300 µM survivin in the absence (*yellow*) and presence (*blue*) of 150 µM peptide VELVNALGQYNDDDDDDDGDDPEEREEKQKDLEDHRDDKE which corresponds to residues 172-211 from EZH2. (**B**) Overlay of 2D [^15^N,^1^H] NMR spectra of 300 µM survivin in the absence (*yellow*) and presence (*dark blue*) of 2.3 mM of the same peptide.

To better mimic the conditions of the microarray experiment and provide kinetic information about the interaction, we performed bio-layer interference (BLI) experiments. We selected three peptides (EZH2_172-211_, EZH2_682-701_, EZH2_722-741_) and one peptide (EED_296-315_) based on the sequences of EZH2 and EED, respectively. The selection of peptides was based on the approximate fluorescence intensity from peptide microarray experiments.

Figure 6 shows BLI interference shift trace profiles associated with the thickness of biolayer because of peptide-survivin interaction. The different peptides interact with survivin (at a low 10 nM concentration) with varied interaction intensity EZH2_172-211_, EZH2_682-701_, EZH2_722-741_ and EED_296-315_ (Figure 6A), whereas sensors without peptides attached peptides consistently show flat or decreasing interference shift over time (Figure 6B). Incubating survivin with the peptide EZH2_172-211_ reached the highest amplitude of interference shift during the association phase of the experiment (Figure 6A) and even at 0.1 nM concentration a qualitatively different interaction curve was obtained (Figure 6C) in contrast to the peptide-free control (Figure 6D). At low survivin concentration aspecific interaction with the sensor is not substantial (Figure 6B and 6D), but above 1 µM survivin concentration it becomes significant (data not shown).

**Figure 6.**
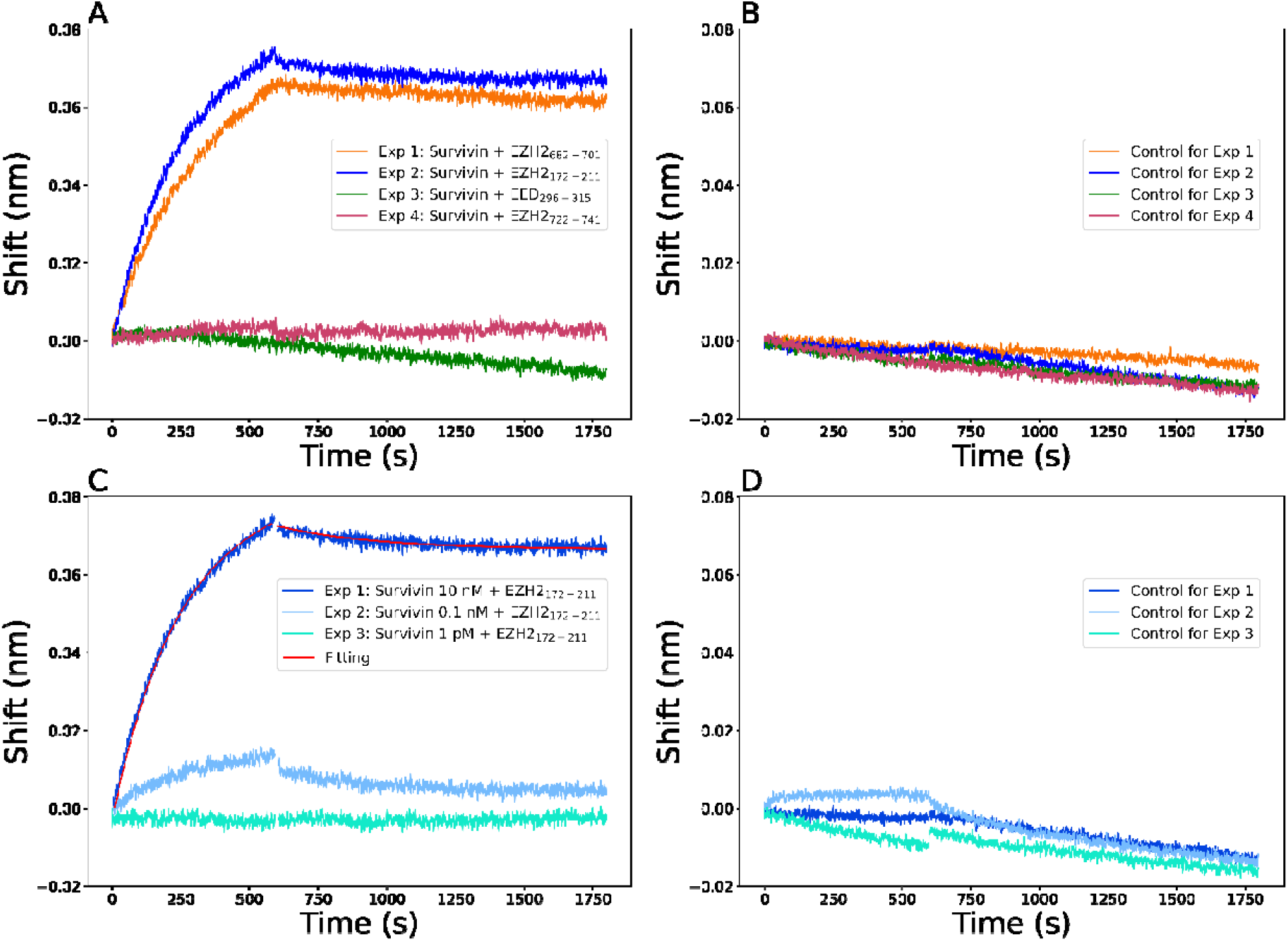
**(A**) Wavelength shift profile from the interaction of 10 nM survivin with peptides EZH2_172-211_, EZH2_682-701_, EZH2_722-741_ and EED_296-315_ (before loading 50 µM each). (**B**) Wavelength shift profile from control biosensors for corresponding experiments in (A). All control biosensors do not contain peptides. (**C**) Wavelength shift profile from the interaction of survivin at different concentrations (10 nM, 0.1 nM, 1 pM) with peptide EZH2_172-211_ (loading 50 µM). Curve fitting (in *red*) of wavelength shift profile (in *blue*) from the interaction of 10 nM survivin with 50 µM (before loading) peptide EZH2_172-211_. The wavelength shift profiles were monitored over association (0-600 s) and dissociation (601-1750 s) periods (**D**) Wavelength shift profile from control biosensors for corresponding experiments in (C). All control biosensors do not contain peptides.

An empirical kinetic model was developed to simulate the association and dissociation phase of the BLI experiment between 10 nM survivin concentration and peptide EZH2_172-211_. This model assumes one and two independent exponential processes for the association and dissociation phases, respectively. The two phases of the experiment were modelled independently with zero wavelength shift as starting point for the association phase and a similarly fixed starting wavelength shift for the dissociation phase. The total modeled amplitudes were constrained to the sum of the independent dissociation exponential processes. While the amplitude was assumed to be unknown for the association phase, the dissociation was expected to reach zero interference shift in the model of the dissociation phase. The modeled progress curves are displayed in Figure 6C. A simple exponential association model was sufficient with an amplitude of 0.081 and a rate constant 3.87 × 10^−3^ s^-1^, respectively. The more complex dissociation model revealed that the fast dissociation process reverted only approximately 7.6 % of the accumulated amplitude (Δλ(0) = 0.071, A_-1_ = 0.005 and A_-2_ = 0.066). The dominant fraction of survivin undergoes very slow dissociation process (*k*_-*1*_ = 2.86 × 10^−3^ s^-1^ and *k*_-*2*_ = 2.40 × 10^−16^ s^-1^). The model predicts that to reach 10% of wavelength shift signal above the pre-association level takes 3.0 × 10^8^ years, which indicates that the interaction is practically irreversible. The dissociation profiles suggest irreversible binding to survivin in all examined peptides above a certain survivin concentration that varies from peptide to peptide. This irreversible binding is scalable and peptide specific. In this light, it is not surprising that in the microarray experiment the relatively harsh washing steps did not remove survivin from the array surface and a wide range of fluorescence intensity was discernible.

## Conclusion

Survivin and PRC2 share functional links and colocalization in the human chromatin. Specific peptides derived from PRC2 subunits attracted survivin at low concentration in a peptide microarray assay and built an irreversible complex. The interacting peptides were frequently originated from important functional regions comprising catalytic, pocket forming, and disease associated amino acid residues. We showed that associating the composition of peptides with colocalization of interaction partners is a promising approach to predict interaction and results in a complex, but reliable pattern. Multiple such peptide regions can be combined into an extremely selective protein-protein targeting system even in the absence of a well-defined 3D structure. This robust targeting mechanism does not require active transport and can move protein molecules against a concentration gradient even in very dilute protein solutions.

## Material and Methods

### Gene expression analysis of CD4+ cells

Blood samples of 24 RA patients were collected at the Rheumatology Clinic, Sahlgrenska Hospital, Gothenburg. All RA patients fulfilled the EULAR/ACR classification criteria (Aletaha 2010 Arthritis Rheum 62, 2569-2581) and gave their written informed consent prior to the blood sampling. The study was approved by the Swedish Ethical Evaluation Board (659-2011) and was performed in accordance with the Declaration of Helsinki. The trial is registered at ClinicalTrials.gov with ID NCT03449589. Human PBMC were isolated using density gradient separation on Lymphoprep (Axis-Shield PoC As, Norway). CD4+ cells were isolated using positive selection (Invitrogen, 11331D), and cultured (1.25×10^6^ cells/ml) in complete RPMI-medium in aCD3 precoated wells (0.5μg/ml, OKT3, Sigma-Aldrich), for 24h.

RNA from the CD4+ cell cultures was prepared using the Norgen Total micro mRNA kit (Norgen, Ontario, Canada). Quality control was done by Bioanalyzer RNA6000 Pico on Agilent2100 (Agilent, St.Clara, CA, USA). Deep sequencing was done by RNAseq (Hiseq2000, Illumina) at the LifeScience Laboratory, Huddinge, Sweden. Raw sequence data were obtained in Bcl-files and converted into fastq text format using the bcl2fastq program from Illumina. Mapping of transcripts was done using Genome UCSC annotation set for hg38 human genome assembly. Analysis was performed using the core Bioconductor packages in R-studio v.3.6.3. Differentially expressed genes were identified using DESeq2 (v.1.26.0) with Benjamini-Hochberg adjustment for multiple testing.

### ChIP seq experiments and analysis

CD4+ cells were isolated from 12 subjects as stated above, stimulated with concanavalin A (0.625μg/ml, Sigma-Aldrich), and LPS (5μg/ml, Sigma-Aldrich) for 72h and pooled in 4 independent samples for chromatin purification. CD4+ cells were cross-linked and lysed according to EpiTect ChIP OneDay kit (Qiagen). After shearing and pre-clearing of chromatin, 1% of the sample was saved as an input. The pre-cleared chromatin was incubated with 2μg anti-Survivin (Tang, Ling, et al (2012) BBRC 421(2) 249-254) (10811, Santa Cruz Biotechnology), anti-H3K27ac (C15410196, Diagenode), anti-H3K27me3 (C15410195, Diagenode) or anti-H3K4m3 (C15411003, Diagenode). The immune complexes were washed, the cross-links reversed, and the DNA purified according to the EpiTect ChIP OneDay kit (Qiagen). Resulting purified DNA was quality controlled using TapeStation (Agilent, Santa Clara, CA, USA). DNA libraries were prepared using ThruPLEX (Rubicon) and sequenced using the Hiseq2000 (Illumina) following the manufacturer’s protocols. Bcl-files were then converted and demultiplexed to fastq using the bcl2fastq program from Illumina.

### ChIP-seq analysis

The fastq sequencing files were mapped to the human reference genome (hg38) using the STAR aligner^42^ with default parameters apart from setting the alignIntronMax flag to 1 for end-to-end mapping. Quality control of the sequenced material was performed by FastQC tool using MultiQC v.0.9dev0 (Babraham Institute, Cambridge, U.K.). Peak calling was performed using the default parameters. Peaks were identified for enrichment of the immunoprecipitation fraction against input (adjusted p<10^−5^) and annotated by HOMER software ^43^ in standard mode to closest transcription start site (TSS). Peaks with overlapping localization by at least 1 nucleotide in several samples were merged and further on referred to as one peak.

### ChIP seq metanalysis

To identify the transcription regulators in vicinity of survivin-ChIP peaks, we performed colocalization analysis of aggregated human ChIPseq datasets of 1034 transcriptional regulators using ReMap database (http://remap.univ-amu.fr/, accessed 15nov2020). Colocalization enrichment analysis was performed using ReMapEnrich R-script (https://github.com/remap-cisreg/ReMapEnrich, accessed 15nov2020). All comparisons were carried out using hg38 human genome assembly. Two-tailed p-values were estimated and normalized by Benjamini-Yekutielli, maximal allowed value of shuffled genomic regions for each dataset (n=15), kept on the same chromosome (shuffling genomic regions parameter byChrom=TRUE). Default fraction of minimal overlap for input and catalogue intervals was set to 10%. Bed interval files of survivin-ChIP peaks with 0, 1 and 10kb flanks were prepared. The dataset with 0kb flanks was compared with the Universe. Statistically significant enrichment of peak overlaps (q-value < 0.05, number >100) were selected.

### Production and purification of survivin

Survivin in pHIS8 ^44^ was expressed in E. coli BL21(DE3) star (Merck). Luria Bertani (LB) medium was used, and expression was induced with 0.5 mM IPTG and incubated for 4 h at 30C. Cells were harvested at 6000 g for 30 minutes and resuspended in lysis buffer containing 50 mM Tris pH 7.5, 300 mM NaCl, 20 mM imidazole, 0.25 mM Pefabloc (Sigma-Aldrich)), 20 mg/ml DNAse (Sigma-Aldrich) and 0.2 mg/ml lysozyme (Sigma-Aldrich). The cells were lysed by sonication with an amplitude of 30 A for 10 minutes (10 s on/30 s off) by a Q700 sonicator (Qsonica). Survivin was purified by immobilized metal ion chromatography (IMAC) with a 5 ml HisTrap FF (GE Healthcare) column followed by overnight dialysis in 50 mM Tris pH 7.5, 150 mM NaCl and 1 mM DTT (survivin SEC buffer) and gel filtration using a Superdex 75 100/300GL column (GE Healthcare).

### Peptide Microarray

A peptide microarray was designed with the subunits of PRC2: EED, EZH2, SUZ12 and JARID2. The protein sequences were divided into peptides of 15 amino acids with an overlap of 10 amino acids. Pre-staining of one of the PEPperCHIP Peptide Microarrays was done with the secondary 6X His Tag Antibody DyLight680 antibody at a dilution of 1:1000 and with monoclonal anti-HA (12CA5)-DyLight800 control antibody at a dilution of 1:1000 to investigate background interactions with the protein-derived peptides that could interfere with the main assays. Subsequent incubation of other peptide microarray copies with survivin at a concentration of 1 μg/ml in incubation buffer was followed by staining with the secondary 6X His Tag Antibody DyLight680 (Rockland Immunochemicals) antibody and the monoclonal anti-HA (12CA5)-DyLight800 control antibody (Rockland Immunochemicals) as well as by read-out at scanning intensities of 7/7 (red/green). HA and His tag control peptides were simultaneously stained as internal quality control to confirm the assay quality and to facilitate grid alignment for data quantification. Read-out was performed with a LI-COR Odyssey Imaging System, while quantification of spot intensities and peptide annotation were done with PepSlide Analyzer. Quantification of spot intensities and peptide annotation were based on the 16-bit gray scale tiff files at scanning intensities of 7/7 that exhibit a higher dynamic range than the 24-bit colorized tiff files shown in Figure S1.

### Classification by machine learning

The scikit-learn python library was used to implement the machine learning process. ^45^ The features of the peptides were the number of atoms that belong to specific atom type categories. For this study, the atoms in the amino acids were assigned according to Table S5. After translating each amino acid in the peptide to atom types, these were accumulated. 5388 peptides from the microarray were divided into equally large training and test sets. The training set was labelled as interacting (fluorescence intensity > 0) or non-interacting (fluorescence intensity = 0). The features were standardized before performing the training. Training was performed with the multi-layer perceptron classifier using the default parameters of scikit-learn. The confusion matrix and prediction accuracy were evaluated by the tools provided by the scikit-learn library.

### BLI experiments on survivin interactions with PRC2 derived peptides

Peptide hits from previous peptide microarray experiments were immobilized using amine reactive second-generation biosensor (AR2G, Fortebio). Ethanolamine was used to quench unbound AR2G surface to prevent non-specific interactions. The biosensor was incubated in MQ water for 15 minutes before use in the Octet instrument (Fortebio) with the following settings: equilibration in water for 60 s and 1000 rpm centrifugation, activation in EDC/s-NHS for 300 s at 1000 rpm, peptide immobilization in immobilization buffer pH 6 (Fortebio) for 900 s and 1000 rpm (50 µM for each dissolved peptide), quenching with 1 M ethanolamine for 300 s and 1000 rpm, baseline measurement in survivin SEC buffer for 600 s and 1000 rpm, association with survivin (100 μM, 1 μM, 10 nM, 0.1 nM, and 1 pM concentrations) for 600 s and 1000 rpm, and dissociation step in survivin SEC buffer for 1200 s and 1000 rpm. The assay was performed at 30 °C. The reference wells and reference biosensors were subtracted by the Octet Data Analysis software.

Δ*λ*(*t*) values were converted to dimensionless quantities by dividing the wavelength shifts with 1 nm prior the analysis. The kinetic modelling was performed with the scipy python package ^46^ using rate equations for the association:

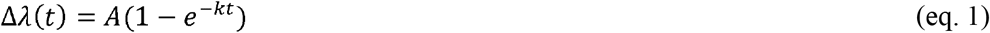

where *A* and *k* are the amplitude and rate constant of the exponential association process, respectively. The rate equation of the dissociation phase was:

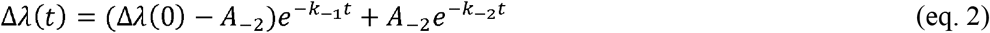

where (Δ*λ*(0) − *λ*_−2_) and *A*_-*2*_ are the amplitudes of the independent exponential dissociation processes with rate constants *k*_-*1*_ and *k*_-*2*_, respectively. The total amplitude is represented by the interference shift at the beginning of the dissociation phase (constant Δ*λ*(0)). The bounds for all parameters were 0 to +∞. The starting values were *A*=1 and *k*=1 s^-1^ for the association and *A*_-*2*_=10^−3^, *k*_-_*1*=10^−4^ s^-1^ and *k*_-*2*_=10^−9^ s^-1^ for the dissociation phase, respectively. The non-linear regression was performed with the default Levenberg-Marquardt algorithm ^47^ of scipy. The residual plots are displayed in Figure S4.

### Isotopic labeling of survivin and NMR titration experiments of survivin by PRC2 derived peptides

^15^N survivin was produced in M9 medium containing ^15^N labeled ammonium chloride (Sigma-Aldrich). Expression was induced with 0.5 mM IPTG and incubated for 20-22 h at 15 °C. Cell harvesting and purification were performed using identical protocol as for unlabeled survivin. The peptides were synthetized and purified to 95% by Biomatik. The peptides were titrated into ^15^N labelled survivin (0.3 mM) at different measure points in the concentration range described in Table 3. Standard HSQC spectra were recorded for ^15^N survivin.

**Table 3.**
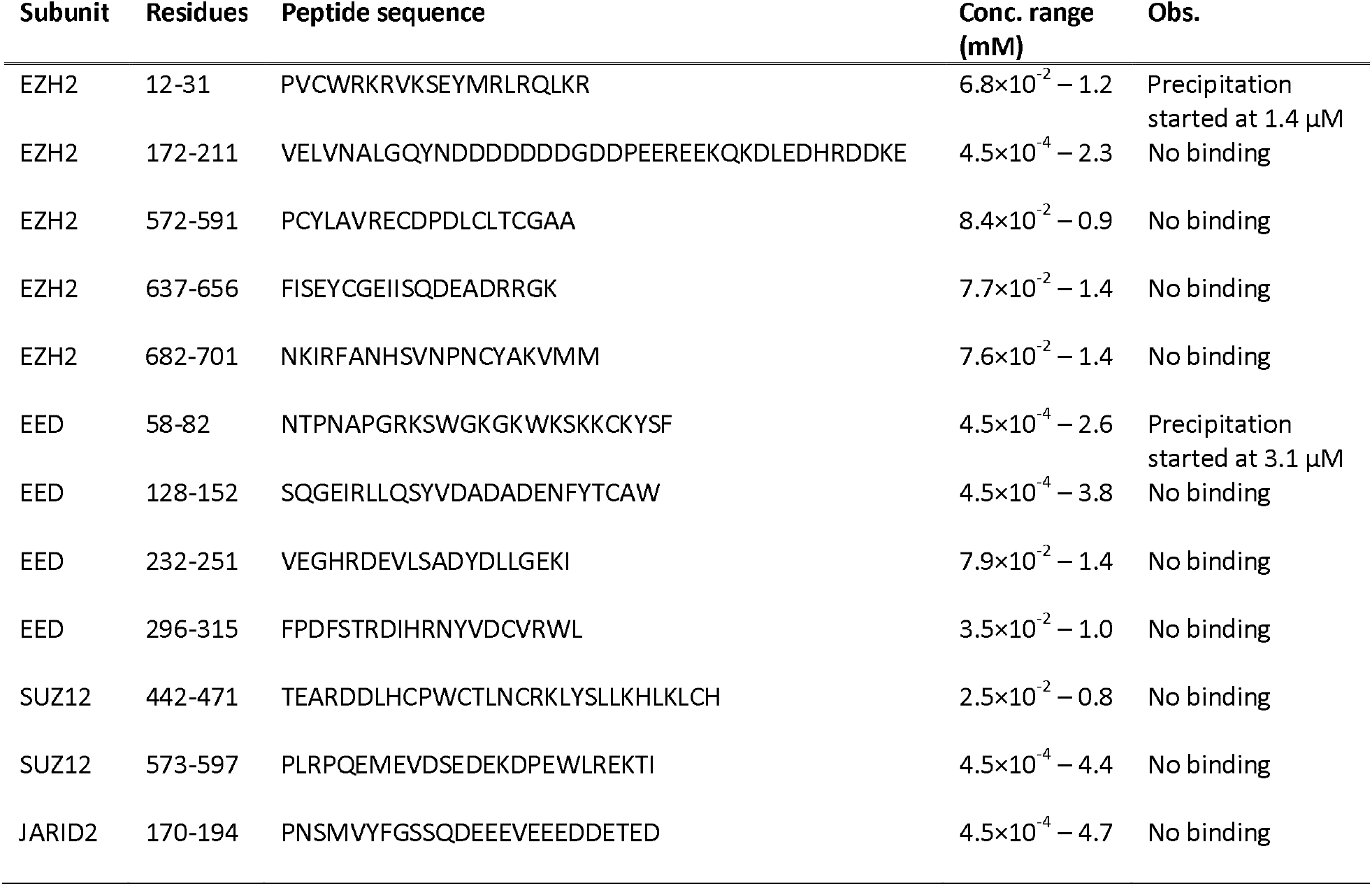
Summary of NMR titration experiments

## Acknowledgements

This work has been funded by grants from the Röntgen-Ångström Cluster Framework of the Swedish Research Council (GK, 2015-06099), the Swedish Research Council (MB, 521-2017-03025), the Swedish Association against Rheumatism (MB, R-566961), the King Gustaf V:s 80-year Foundation (MB), the Regional agreement on medical training and clinical research between the Western Götaland county council and the University of Gothenburg (MB, ALFGBG-717681). This project has received funding from the European Union’s Horizon 2020 research and innovation programme under grant agreement No 964203.

