## Supplementary tables and figures for "Survivin prevents the Polycomb Repressor Complex 2 from methylating Histone 3 lysine 27"

**Table S1** Annotated features in the EZH2 peptides interacting with survivin.

| EZH2 | Sequence | Fluorescence intensity (au, 0-64193) | Comments |
| --- | --- | --- | --- |
| S21 | KSEKGPVCWRKRVK**S** | 4969 | Phosphorylation, methyltransferase activity^1^ |
|  | PVCWRKRVK**S**EYMRL | 679 |  |
|  | KRVK**S**EYMRLRQLKR | 19209 |  |
| P132 & Y133 | VEDETVLHNIPYMGD | 1640 | Weaver syndrome, impairs methyltransferase^2^ |
|  | VLHNIPYMGDEVLDQ | 889 |  |
|  | PYMGDEVLDQDGTFI | 1993 |  |
| C588 | PDLCLT**C**GAADHWDS | 1287 | Impairs methyltransferase^3^ |
|  | T**C**GAADHWDSKNVSC | 682 |  |
| Y641 | VQKNEFISE**Y**CGEII | 1037 | active site |
|  | FISE**Y**CGEIISQDEA | 2238 |  |
| Y726 | AKRAIQTGEELFFD**Y** | 17722 | active site |
|  | QTGEELFFD**Y**RYSQA | 1545 |  |
| Y726 & Y736 | LFFD**Y**RYSQADALK**Y** | 957 | active site & Weaver syndrome^4^ |
| Y736 & E740 | RYSQADALK**Y**VGI**E**R | 552 | Weaver syndrome^4^ |
|  | DALK**Y**VGI**E**REMEIP | 82 |  |

**Table S2** Annotated features in the EED interacting peptides

| EED | Sequence | Fluorescence intensity (au, 0-64193) | Comments |
| --- | --- | --- | --- |
| K70-K79 | NTPNAPGRKSWGKGK | 2749 | DNA binding^5^ |
|  | PGRKSWGKGKWKSKK | 13033 |  |
|  | WGKGKWKSKKCKYSF | 5143 |  |
|  | WKSKKCKYSFKCVNS | 112 |  |
| Y148 | YVDADADENF**Y**TCAW | 5116 | aromatic cage^6^ |
|  | ADENF**Y**TCAWTYDSN | 1628 |  |
| R236 | DTLVAIFGGVEGH**R**D | 539 | Weaver syndrome |
|  | IFGGVEGH**R**DEVLSA | 93 |  |
|  | EGH**R**DEVLSADYDLL | 5179 |  |
| R302 & Y308 | DFST**R**DIHRN**Y**VDCV | 579 | Weaver syndrome, alanine binding pocket |
|  | DIHRN**Y**VDCVRWLGD | 1492 |  |
| C324 | LILSKS**C**ENAIVCWK | 120 | alanine binding pocket ^6^ |
| W364 & Y365 | LGRFDYSQCDI**WY**MR | 678 | aromatic cage^6^ |
|  | YSQCDI**WY**MRFSMDF | 1624 |  |

**Table S3** Annotated features in the SUZ12 interacting peptides.

| SUZ12 | Sequence | Fluorescence intensity (au, 0-64193) | Comments |
| --- | --- | --- | --- |
| Zinc finger (448 – 471) | EARDDLHCPWCTLNC | 1389 |  |
|  | CTLNCRKLYSLLKHL | 338 |  |
|  | LLKHLKLCHSRFIFN | 212 |  |
| E610 | TQIEEFSDVNEGEKE | 2471 | Weaver syndrome^7^ |
|  | FSDVNEGEKEVMKLW | 646 |  |

**Table S4** Annotated features in the JARID2 interacting peptides

| JARID2 | Sequence | Fluorescence intensity (au, 0-64193) | Comments |
| --- | --- | --- | --- |
| ARID domain | RWGPNVQRLACIKKH | 3046 |  |
|  | VQRLACIKKHLKSQG | 165 |  |
|  | LKSQGITMDELPLIG | 56 |  |
|  | ITMDELPLIGGCELD | 6829 |  |
|  | LPLIGGCELDLACFF | 2771 |  |
|  | GCELDLACFFRLINE | 1943 |  |
|  | AQDRLAKLQEAYCQY | 730 |  |
|  | AKLQEAYCQYLLSYD | 7908 |  |
|  | AYCQYLLSYDSLSPE | 1935 |  |
|  | LLSYDSLSPEEHRRL | 316 |  |

**Table S5** Amino acid – atom type conversion table used for the peptide analysis.

| aa | CA-Gly | Pro-MC | Carboxyl | Amide | His | Trp | Phe-Tyr | OH-Tyr | CH2 | CH | CH3 | OH | SH | S | NH3 | Arg | MC |
| --- | --- | --- | --- | --- | --- | --- | --- | --- | --- | --- | --- | --- | --- | --- | --- | --- | --- |
| A | 0 | 0 | 0 | 0 | 0 | 0 | 0 | 0 | 0 | 0 | 1 | 0 | 0 | 0 | 0 | 0 | 4 |
| C | 0 | 0 | 0 | 0 | 0 | 0 | 0 | 0 | 1 | 0 | 0 | 0 | 1 | 0 | 0 | 0 | 4 |
| D | 0 | 0 | 3 | 0 | 0 | 0 | 0 | 0 | 1 | 0 | 0 | 0 | 0 | 0 | 0 | 0 | 4 |
| E | 0 | 0 | 3 | 0 | 0 | 0 | 0 | 0 | 2 | 0 | 0 | 0 | 0 | 0 | 0 | 0 | 4 |
| F | 0 | 0 | 0 | 0 | 0 | 0 | 6 | 0 | 1 | 0 | 0 | 0 | 0 | 0 | 0 | 0 | 4 |
| G | 1 | 0 | 0 | 0 | 0 | 0 | 0 | 0 | 0 | 0 | 0 | 0 | 0 | 0 | 0 | 0 | 3 |
| H | 0 | 0 | 0 | 0 | 5 | 0 | 0 | 0 | 1 | 0 | 0 | 0 | 0 | 0 | 0 | 0 | 4 |
| I | 0 | 0 | 0 | 0 | 0 | 0 | 0 | 0 | 1 | 1 | 2 | 0 | 0 | 0 | 0 | 0 | 4 |
| K | 0 | 0 | 0 | 0 | 0 | 0 | 0 | 0 | 4 | 0 | 0 | 0 | 0 | 0 | 1 | 0 | 4 |
| L | 0 | 0 | 0 | 0 | 0 | 0 | 0 | 0 | 1 | 1 | 2 | 0 | 0 | 0 | 0 | 0 | 4 |
| M | 0 | 0 | 0 | 0 | 0 | 0 | 0 | 0 | 2 | 0 | 1 | 0 | 0 | 1 | 0 | 0 | 4 |
| N | 0 | 0 | 0 | 3 | 0 | 0 | 0 | 0 | 1 | 0 | 0 | 0 | 0 | 0 | 0 | 0 | 4 |
| P | 0 | 2 | 0 | 0 | 0 | 0 | 0 | 0 | 3 | 0 | 0 | 0 | 0 | 0 | 0 | 0 | 2 |
| Q | 0 | 0 | 0 | 3 | 0 | 0 | 0 | 0 | 2 | 0 | 0 | 0 | 0 | 0 | 0 | 0 | 4 |
| R | 0 | 0 | 0 | 0 | 0 | 0 | 0 | 0 | 3 | 0 | 0 | 0 | 0 | 0 | 0 | 4 | 4 |
| S | 0 | 0 | 0 | 0 | 0 | 0 | 0 | 0 | 1 | 0 | 0 | 1 | 0 | 0 | 0 | 0 | 4 |
| T | 0 | 0 | 0 | 0 | 0 | 0 | 0 | 0 | 0 | 1 | 1 | 1 | 0 | 0 | 0 | 0 | 4 |
| Y | 0 | 0 | 0 | 0 | 0 | 0 | 6 | 1 | 1 | 0 | 0 | 0 | 0 | 0 | 0 | 0 | 4 |
| V | 0 | 0 | 0 | 0 | 0 | 0 | 0 | 0 | 0 | 1 | 2 | 0 | 0 | 0 | 0 | 0 | 4 |
| W | 0 | 0 | 0 | 0 | 0 | 9 | 0 | 0 | 1 | 0 | 0 | 0 | 0 | 0 | 0 | 0 | 4 |

A
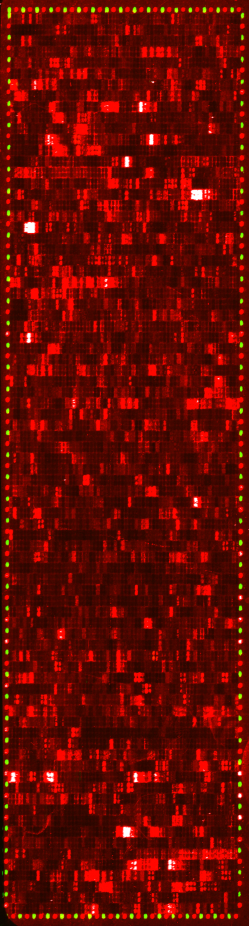
 B
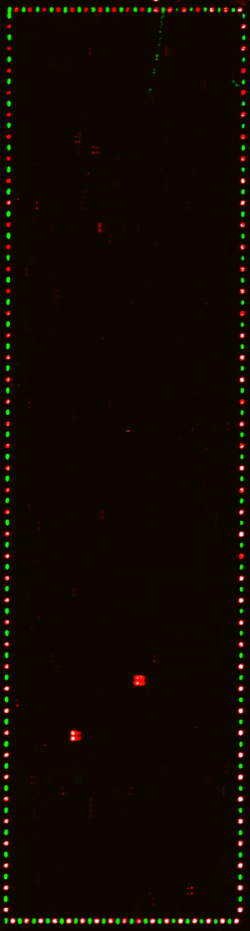

**Figure S1** Raw fluorescence scans of the peptide microarray (A) incubated with 1 µg/mL survivin and labelled anti-His tag antibody (B) only labelled anti-His tag antibody.

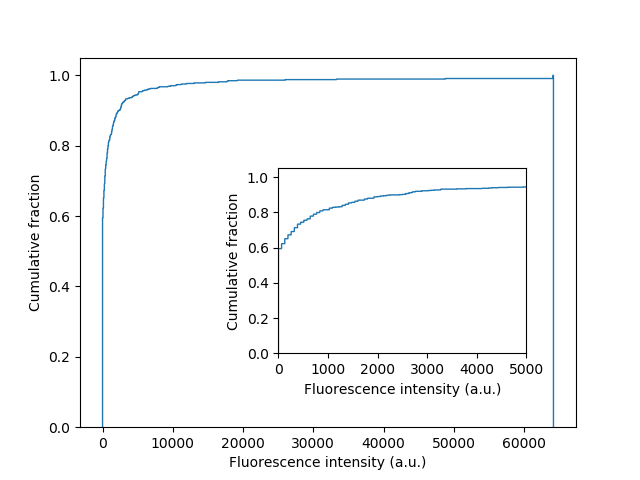

**Figure S2** Cumulative fraction of fluorescence intensity less than a certain value in the peptide microarray experiment.

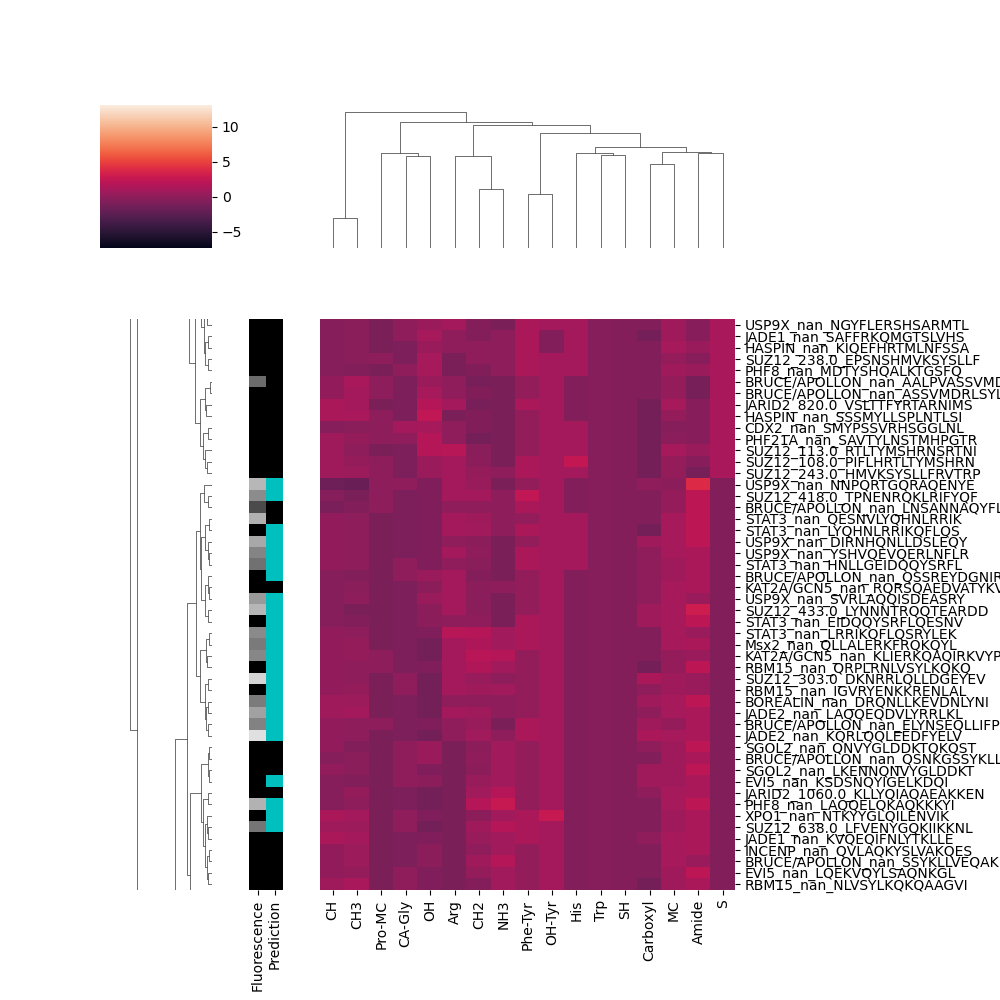

**Figure S3** Detailed view on connection between survivin interaction with peptides and the peptide composition. A zoomed in region of the cluster where individual peptides are discernible. The heat map shows light/dark high/low abundance of atom types, respectively (z-score). The grayscale bar indicates the logarithm of fluorescence intensity of the peptide in the survivin peptide microarray experiment (*black* zero intensity, *white* highest level of intensity). The prediction bar shows the success of the machine learning prediction using the atom type abundance as features (*black* and *cyan* colors mark predicted non-interacting and interacting peptides, respectively).

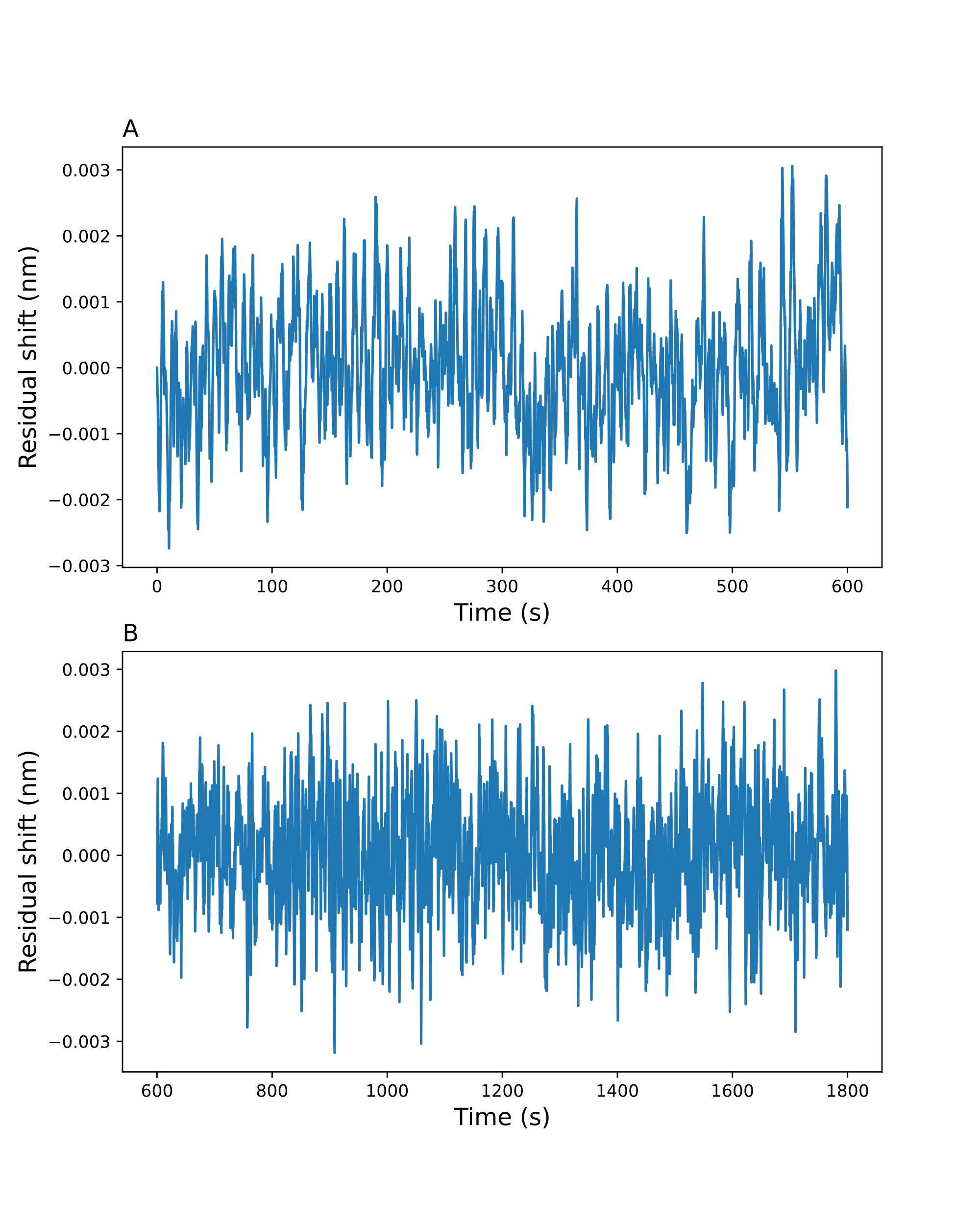

**Figure S4** Residual plots of (A) association and (B) dissociation phase.
